# CXCL17 activates three MAS-related G protein-coupled receptors independently of its conserved C-terminal fragment

**DOI:** 10.1101/2025.05.20.655027

**Authors:** Wen-Feng Hu, Juan-Juan Wang, Jie Yu, Ya-Li Liu, Zeng-Guang Xu, Zhan-Yun Guo

## Abstract

C-X-C motif chemokine ligand 17 (CXCL17) is a chemoattractant whose receptor remains controversial. While recent studies identified CXCL17 as an agonist of G protein-coupled receptor 25 (GPR25), it was also reported to activate MAS-related GPR family member X2 (MRGPRX2), a member of MAS-related G protein-coupled receptors (MRGPRs), though this finding has not yet been reproduced by other laboratories. In this study, we confirmed that micromolar concentrations of human CXCL17 activate human MRGPRX2 in transfected human embryonic kidney (HEK) 293T cells using a NanoLuc Binary Technology (NanoBiT)-based β-arrestin recruitment assay. We further demonstrated that human CXCL17 also activates MRGPRX1 and MAS1 among 10 human MRGPRs in the same assay. CXCL17 could also induce chemotactic movement of transfected HEK293T cells expressing MRGPRX2, MRGPRX1, or MAS1. However, removal of C-terminal residues from CXCL17 did not affect its activation of these three MRGPRs, even though this region is essential for GPR25 activation. These results suggest that CXCL17 activates MRGPRX2, MRGPRX1, and MAS1 through a mechanism distinct from GPR25 activation. Further investigation is needed to determine whether these MRGPRs mediate the *in vivo* functions of CXCL17.

## 1. Introduction

C-X-C motif chemokine ligand 17 (CXCL17), previously referred to as dendritic cell and monocyte chemokine-like protein (DMC) or VEGF-correlated chemokine 1 (VCC-1), was first identified in 2006 [1,2]. It functions as a chemoattractant for leukocytes, including monocytes, macrophages, and dendritic cells [1,3□14], and is also implicated in angiogenesis and tumor progression [2,15□22]. Given these roles, it is presumed that CXCL17 exerts its functions through specific plasma membrane receptor(s). Although CXCL17 was initially proposed as an agonist of the orphan G protein-coupled receptor 35 (GPR35) [23], this pairing cannot be reproduced by other laboratories [22□26]. More recently, CXCL17 has been reported to act on the MAS-related G protein-coupled receptor X2 (MRGPRX2), to function as an allosteric regulator of C-X-C motif chemokine receptor 4 (CXCR4), or to interact with extracellular glycosaminoglycans [26□28]. Most recently, both Ocón’s group and our group identified CXCL17 as an agonist of the rarely studied orphan G protein-coupled receptor 25 (GPR25) [29,30].

MRGPRX2 belongs to the MAS-related G protein-coupled receptor (MRGPR) family, which includes 10 human paralogs: MAS1, MAS1L, MRGPRD, MRGPRE, MRGPRF, MRGPRG, MRGPRX1, MRGPRX2, MRGPRX3, and MRGPRX4 (Table S1). These receptors share substantial sequence similarity (Fig. S1) and are linked to immune and sensory functions [31□34]. However, all remain officially classified as orphan receptors. According to the gene database at the National Center for Biotechnology Information (NCBI), MRGPRX2 orthologs are widely distributed in eutherians and share significant sequence similarity (Fig. S2). However, MRGPRX2 orthologs are absent from lower mammals and other lower vertebrates. MRGPRX2 is primarily expressed in mast cells and plays a role in allergy and itching, making it a potential therapeutic target [34□38]. MRGPRX2 has a broad ligand spectrum and can be activated by various peptides and compounds at micromolar concentrations, but its endogenous ligand remains unknown [34].

Independent confirmation is essential for establishing ligand□receptor pairings [39,40]. This study aimed to investigate whether human CXCL17 activates human MRGPRX2, as well as other human MRGPRs. Using the NanoLuc Binary Technology (NanoBiT)-based β-arrestin recruitment assay in transfected human embryonic kidney (HEK) 293T cells, we observed that micromolar concentrations of recombinant human CXCL17 not only activated MRGPRX2, consistent with the previous report [27], but also activated MRGPRX1 and MAS1 among 10 human MRGPRs. Furthermore, CXCL17-mediated activation of these MRGPRs induced chemotactic movement of the transfected HEK293T cells. Interestingly, the conserved C-terminal fragment of CXCL17 was dispensable for activating the three MRGPRs, despite being crucial for activating GPR25. These findings suggest that CXCL17 activates these MRGPRs through a distinct mechanism than it uses to activate GPR25. Further research is necessary to determine whether these MRGPRs mediate the *in vivo* functions of CXCL17.

## 2. Experimental methods

### 2.1. Preparation of the recombinant human CXCL17s

The expression constructs for wild-type (WT) CXCL17 and the C-terminally truncated [desC3]CXCL17 were generated in our recent study [30]. The expression construct for [desC8]CXCL17 was generated via the QuikChange approach using pET/6×His-[desC3]CXCL17 as the mutagenesis template. Thereafter, the N-terminally 6×His-tagged WT or mutant CXCL17s were overexpressed in *Escherichia coli* as inclusion bodies and purified by immobilized metal ion affinity chromatography according to our previous procedure [30]. After *in vitro* refolding and removal of the N-terminal tag via enterokinase cleavage, mature CXCL17 proteins were purified by high performance liquid chromatography (HPLC) using a C_18_ reverse-phase column (Zorbax 300SB-C18, 9.4 × 250 mm, Agilent Technologies, Santa Clara, CA, USA). After lyophilization, the mature WT or mutant CXCL17s were dissolved in 1.0 mM aqueous hydrochloride (pH 3.0), quantified by ultra-violet absorbance at 280 nm using the extinction coefficient of *ε*_280 nm_ = 11000 M^-1^ cm^-1^, and analyzed by sodium dodecyl sulfate-polyacrylamide gel electrophoresis (SDS-PAGE).

### 2.2. Generation of the expression constructs for human MRGPRs

Information of 10 human MRGPRs was retrieved from the gene database of NCBI (Table S1) and their coding regions were amplified by polymerase chain reaction (PCR) using human genomic DNA (extracted from HEK293T cells) as template and synthetic oligoes as primers (Table S2). After cleavage with appropriate restriction enzymes (Table S2), the amplified DNA fragments were ligated into a pcDNA6 vector and confirmed by DNA sequencing. To generate the constructs for NanoBiT-based β-arrestin recruitment assay, their coding regions were PCR amplified using appropriate primers and ligated into a pTRE3G-BI/SmBiT-ARRB2 vector via Gibson assembly (Table S2). The resultant construct pTRE3G-BI/MRGPR-LgBiT:SmBiT-ARRB2 coexpresses a C-terminally large NanoLuc fragment (LgBiT)-fused human MRGPR (MRGPR-LgBiT) and an N-terminally low-affinity complementation tag (SmBiT)-fused human β-arrestin 2 (SmBiT-ARRB2) under control of a doxycycline (Dox)-response bidirectional promoter. Untagged MAS1, MRGPRX1, and MRGPRX2 were also subcloned from the pcDNA6 vector into a Dox-response PB-TRE vector for selection stably transfected HEK293T cells (Table S2).

### 2.3. β*-arrestin recruitment assays*

β-arrestin recruitment assays of these MRGPRs were conducted according to our previous procedure for other GPCRs [30,41,42]. Briefly, the construct pTRE3G-BI/MRGPR-LgBiT:SmBiT-ARRB2 was transfected into HEK293T cells together with the expression control vector pCMV-TRE3G (Clontech, Mountain View, CA, USA) using the transfection reagent Lipo8000 (Beyotime Technology, Shanghai, China). The next day, the transfected cells were trypsinized, suspended in the induction medium (complete DMEM medium plus 1.0□10 ng/mL of Dox), seeded into white opaque 96-well plates, and cultured at 37 °C for ~24 h to ~90% confluence. To conduct the assay, the medium was removed and pre-warmed activation solution (serum-free DMEM plus 1% bovine serum albumin) containing NanoLuc substrate was added (40 μL/well, containing 1.0 μL of NanoLuc substrate stock from Promega, Madison, WI, USA), and bioluminescence data were immediately collected for ~4 min on a SpectraMax iD3 plate reader (Molecular Devices, Sunnyvale, CA, USA). Subsequently, WT or mutant CXCL17 (diluted in the activation solution) was added (10 μL/well), and bioluminescence data were continuously collected for ~10 min. The measured absolute signals were corrected for inter well variability by forcing all curves after addition of NanoLuc substrate (without ligand) to same level and plotted using the SigmaPlot 10.0 software (SYSTAT software, Chicago, IL, USA).

### 2.4. Transwell chemotaxis assay

The transwell chemotaxis assay was conducted on transfected HEK293T cells according to our recent procedure [30]. The expression construct for the untagged MRGPRX2, MRGPRX1, or MAL1 in the PB-TRE vector was transfected into HEK293T cells together with the Super PiggyBac Transposase expression vector (System Biosciences, CA, USA) using the Lipo8000 transfection reagent (Beyotime Technology). From the next day, the transfected cells were treated with 200 μg/mL of hygromycin B (Beyotime Technology) for about one month, and the resultant cell pool was used for transwell chemotaxis assays conducted in 24-well transwell apparatuses (Corning, NY, USA). Before the assay, the stably transfected HEK293T cells were cultured in the induction medium (complete medium plus 3.0 ng/mL of Dox) for ~24 h. Thereafter, the cells were trypsinized, washed, suspended in serum-free DMEM at the density of ~5×10^5^ cells/mL, and seeded into polyethylene terephthalate membrane (8 μm pore size)-coated permeable inserts (150 μL/well). The inserts were put into the lower plate chambers containing 500 μL of chemotactic agent (serum-free DMEM plus indicated concentrations of WT CXCL17). After cultured at 37 °C for ~6.5 h, solution in the inserts were removed and cells on the upper face of the permeable membrane were wiped off using cotton swaps. Thereafter, cells on the lower face of the permeable membrane were fixed with 4% paraformaldehyde solution, stained with crystal violet staining solution (Beyotime Technology), and observed under an Olympus APX100 microscope (Tokyo, Japan).

## 3. Results

### 3.1. Human CXCL17 can activate three human MRGPRs

To test whether CXCL17 can activate MRGPRX2 or other MRGPRs, we employed the NanoBiT-based β-arrestin recruitment assay. This assay is unlikely to be affected by endogenously expressed CXCL17 receptors because they lack the LgBiT fusion. To set up this assay, 10 human MRGPRs were genetically fused with an inactive LgBiT at their intracellular C-terminus and coexpressed with an N-terminally SmBiT-fused human β-arrestin 2 (SmBiT-ARRB2) in transfected HEK293T cells under the control of a doxycycline (Dox)-response bidirectional promoter. If an MRGPR-LgBiT could be activated by CXCL17, it would recruit the coexpressed SmBiT-ARRB2. This proximity effect would induce complementation of the β-arrestin-fused SmBiT with the receptor-fused LgBiT, and restore luciferase activity.

After the transiently transfected HEK293T cells were induced to coexpress MRGPRX2-LgBiT and SmBiT-ARRB2, low bioluminescence was detected following the addition of NanoLuc substrate (Fig. 1A), indicating background complementation of the coexpressed MRGPRX2-LgBiT and SmBiT-ARRB2. Subsequent addition of recombinant human CXCL17 resulted in a rapid, dose-dependent increase in measured bioluminescence (Fig. 1A). Concentrations as low as 0.1 μM of CXCL17 caused a significant activation effect, suggesting that CXCL17 indeed activates MRGPRX2 in the micromolar range, as previously reported [27].

**Fig. 1.**
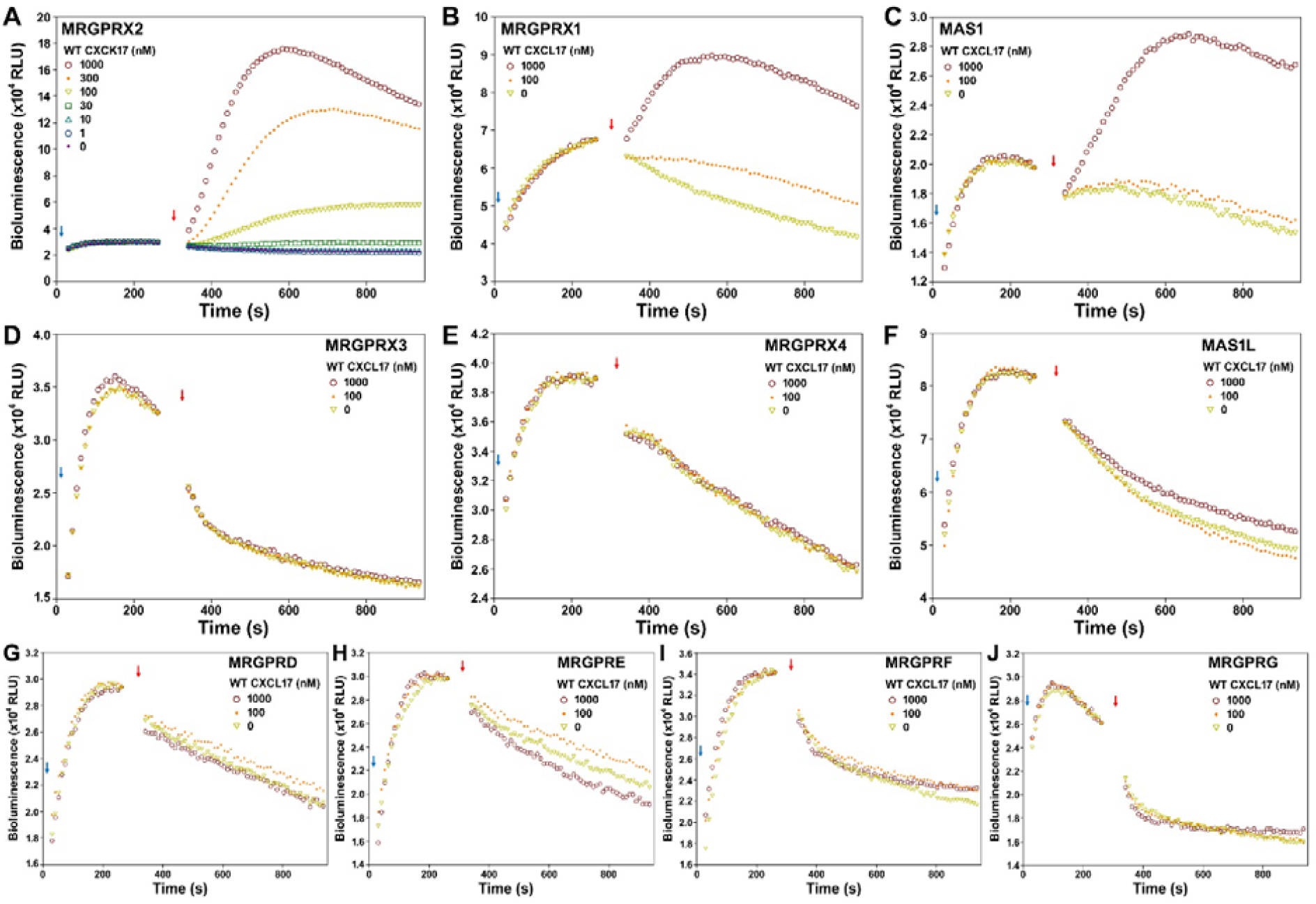
Effect of human CXCL17 on human MRGPRs in the NanoBiT-based β-arrestin recruitment assays. A typical bioluminescence change after sequential addition of NanoLuc substrate and WT CXCL17 to living HEK293T cells coexpressing SmBiT-ARRB2 and MRGPRX2-LgBiT (**A**), MRGPRX1-LgBiT (**B**), MAS1-LgBiT (**C**), MRGPRX3-LgBiT (**D**), MRGPRX4-LgBiT (**E**), MAS1L-LgBiT (**F**), MRGPRD-LgBiT (**G**), MRGPRE-LgBiT (**H**), MRGPRF-LgBiT (**I**), or MRGPRG-LgBiT (**J**). In these panels, blue arrows indicate the addition of NanoLuc substrate and red arrows indicate the addition of WT CXCL17. The panels are representatives of two independent experiments.

In HEK293T cells coexpressing MRGPRX1-LgBiT and SmBiT-ARRB2, the addition of human CXCL17 also caused a dose-dependent increase in bioluminescence (Fig. 1B). In HEK293T cells coexpressing MAS1-LgBiT and SmBiT-ARRB2, 1.0 μM of human CXCL17 induced a significant increase in bioluminescence (Fig. 1C). However, in HEK293T cells coexpressing other MRGPR-LgBiT and SmBiT-ARRB2, up to 1.0 μM of recombinant human CXCL17 had no detectable activation effect (Fig. 1D□J). Thus, micromolar concentrations of human CXCL17 could activate three human MRGPRs, namely MRGPRX2, MRGPX1, and MAS1, in our β-arrestin recruitment assay.

### 3.2. CXCL17 activates the three MRGPRs independently of its conserved C-terminal fragment

The conserved C-terminal fragment is essential for CXCL17 to activate GPR25 [29,30]. We investigated whether it is also responsible for CXCL17 activating MRGPRs. To address this, we first used a truncated CXCL17, [desC3]CXCL17, lacking three C-terminal residues, which abolished its activation effect on human GPR25 [30]. Surprisingly, [desC3]CXCL17 activated human MRGPRX2, MRGPRX1, and MAS1 as efficiently as wild-type (WT) CXCL17 in the β-arrestin recruitment assay (Fig. 2), indicating that the three C-terminal residues are dispensable for CXCL17 activating these MRGPRs. Removal of eight C-terminal residues also did not affect activation, as the truncated [desC8]CXCL17 activated these MRGPRs as efficiently as WT CXCL17 (Fig. 2). These results suggest that CXCL17 activates the three MRGPRs independently of its conserved C-terminal fragment, implying the use of different mechanisms to activate GPR25 and these three MRGPRs.

**Fig. 2.**
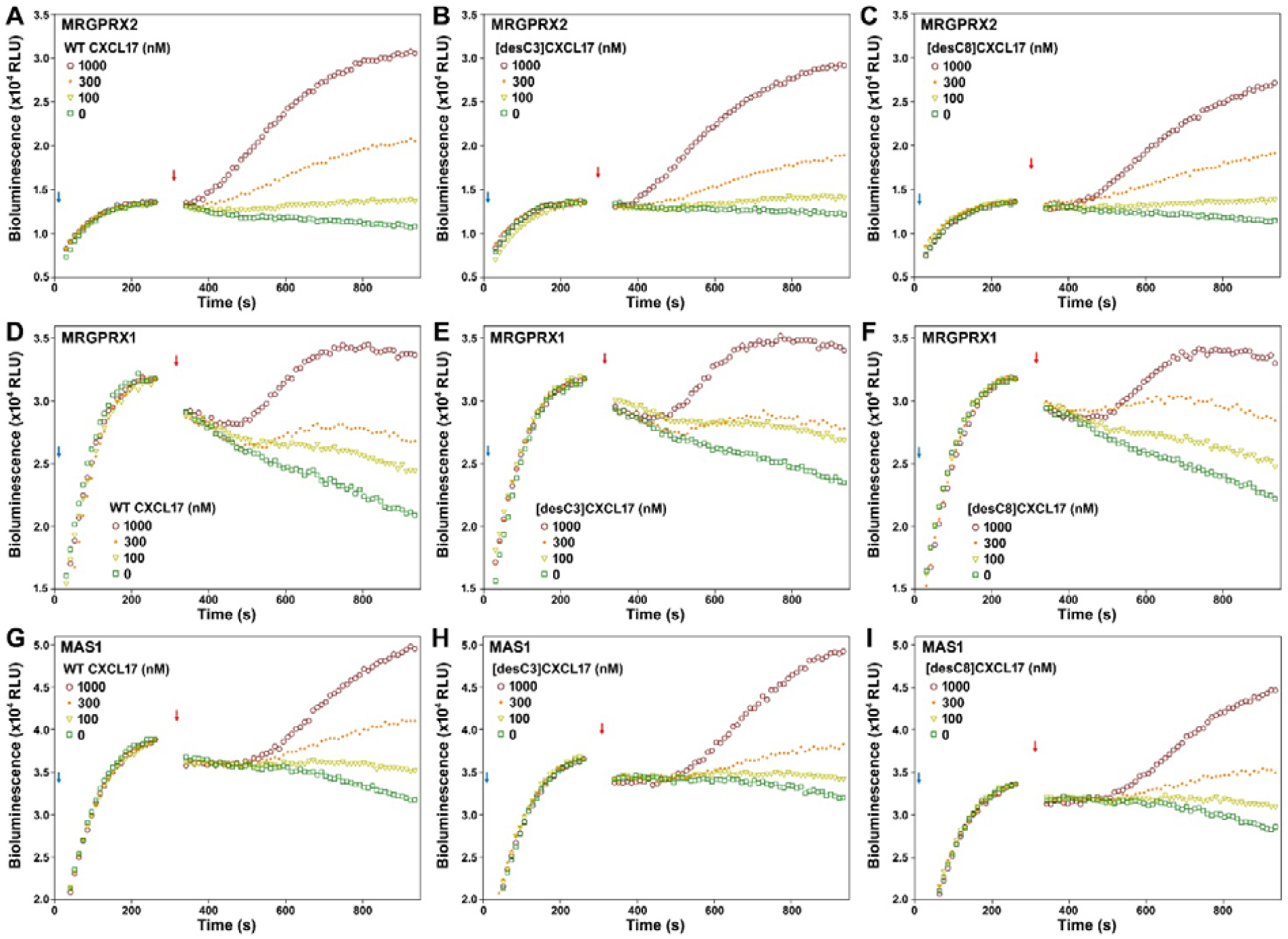
Effect of WT CXCL17 or C-terminally truncated mutants on some human MRGPRs in the NanoBiT-based β-arrestin recruitment assay. A typical bioluminescence change after sequential addition of NanoLuc substrate and WT CXCL17 (**A**,**D**,**G**), [desC3]CXCL17 (**B**,**E**,**H**), or [desC8]CXCL17 (**C**,**F**,**I**) to living HEK293T cells coexpressing SmBiT-ARRB2 and MRGPRX2-LgBiT (**A**,**B**,**C**), MRGPRX1-LgBiT (**D**,**E**,**F**), or MAS1-LgBiT (**G**,**H**,**I**). In these panels, blue arrows indicate the addition of NanoLuc substrate and red arrows indicate the addition of WT or mutant CXCL17. The panels are representatives of two independent experiments.

### 3.3. CXCL17 induces chemotactic movement of HEK293T cells expressing the three MRGPRs

Since CXCL17 functions as a chemoattractant, we examined whether it could induce chemotactic movement of transfected HEK293T cells expressing MRGPRX2, MRGPRX1, or MAS1. Our previous study demonstrated that up to 1.0 μM of WT CXCL17 failed to induce migration of untransfected HEK293T cells in the transwell chemotaxis assay, suggesting that HEK293T cells do not express endogenous CXCL17 receptors. When stably transfected HEK293T cells were induced to express human MRGPRX2, recombinant human CXCL17 induced cell migration in a dose-dependent manner (Fig. 3A), with significant effects observed at concentrations as low as 30 nM. Similarly, recombinant CXCL17 significantly affected cell migration when stably transfected HEK293T cells were induced to express human MRGPRX1 or MAS1 (Fig. 3B, C), although the effect was slightly less potent than that observed with cells expressing MRGPRX2 (Fig. 3A). Collectively, these findings indicate that CXCL17 can induce chemotactic movement of transfected HEK293T cells via activation of MRGPRX2, MRGPRX1, or MAS1.

**Fig. 3.**
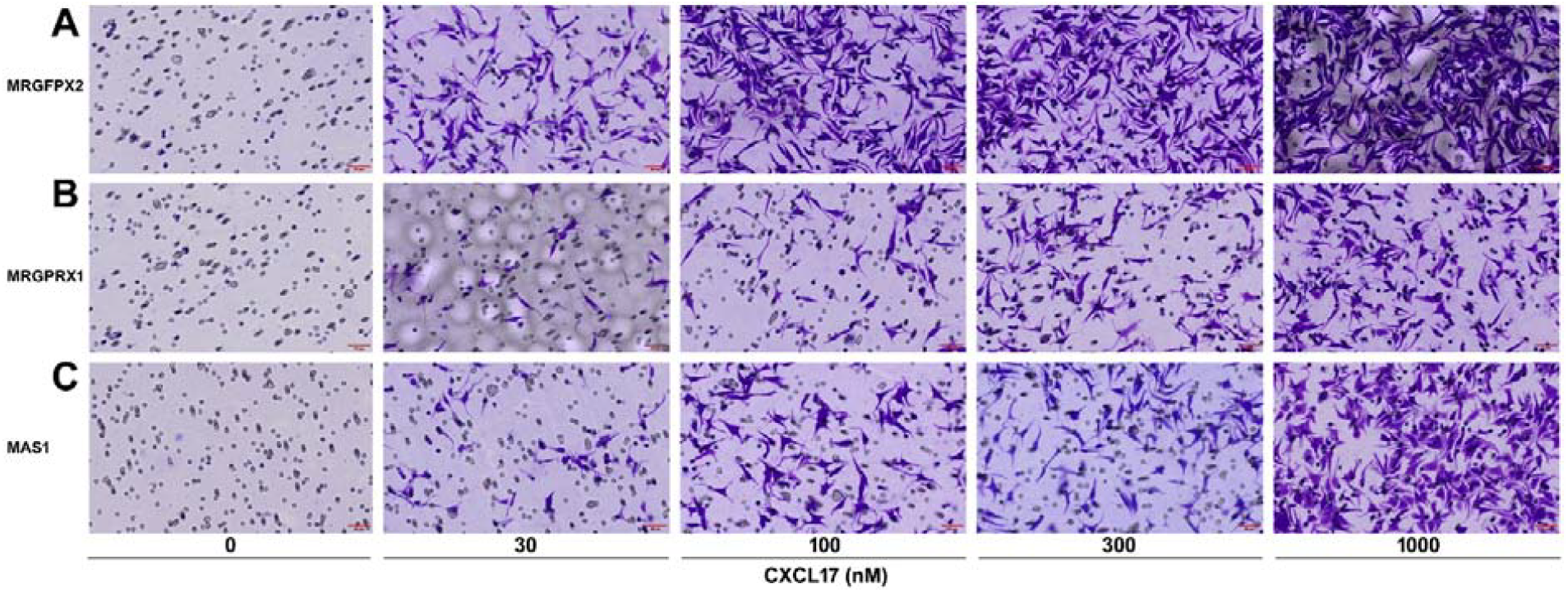
Effect of human CXCL17 on chemotactic movement of stably transfected HEK293T cells expressing the indicated human MRGPR. The stably transfected HEK293T cells were induced to express human MRGPRX2 (**A**), MRGPRX1 (**B**), or MAS1 (**C**) and then seeded into the permeable membrane-coated inserts, and serum-free DMEM containing indicated concentrations of WT human CXCL17 was added to the lower plate chambers. After the assay, cells on the upper face of the permeable membrane were wiped off, and cells on the lower face of the permeable membrane were fixed, stained, and observed under a microscope. These panels are representatives of two independent experiments.

## 4. Discussion

Using the NanoBiT-based β-arrestin recruitment assay and chemotaxis assay conducted on transfected HEK293T cells, we not only confirmed that human CXCL17 can activate human MRGPRX2, as recently reported by Ding et al. [27], but also demonstrated that human CXCL17 can activate two other members of human MRGPRs, namely MRGPRX1 and MAS1, albeit with slightly lower potency. Thus, three members of human MRGPRs can be activated by human CXCL17, and this activation can induce chemotactic movement of the transfected HEK293T cells. Further studies are needed to examine whether the *in vivo* functions of CXCL17 are mediated by these MRGPRs.

Recent studies have shown that CXCL17 activates GPR25 via its C-terminal fragment [29,30], which aligns with the observation that some C-terminal residues are highly conserved in CXCL17 orthologs across different species. However, our present study demonstrates that the conserved C-terminal fragment is dispensable for CXCL17 activating these MRGPRs, suggesting that CXCL17 employs different mechanisms to activate GPR25 and these three MRGPRs. Future research should focus on identifying the specific residues responsible for CXCL17 activating these MRGPRs through mutational analysis.

Both MRGPRX2 and MRGPRX1 are implicated in allergy and itching and can be activated by a variety of peptides and compounds [31□38, 43□46]. MAS1, also known as MAS or MAS receptor (MasR) in the literatures, can mediate the function of angiotensin-(1-7) and plays a role in hypotension and smooth muscle relaxation [47□50]. According to data from the Human Protein Atlas (https://www.proteinatlas.org), human MAS1 is expressed by various cell types, including certain immune cells, such as B-cells, T-cells, and Langerhans cells, implying that MAS1 likely plays a role in immunity. Future studies should clarify whether MRGPRX2, MRGPRX1, and MAS1 mediate the *in vivo* function of CXCL17.

## CRediT authorship contribution statement

Wen-Feng Hu, Jian-Juan Wang and Jie Yue performed the experiments; Ya-Li Liu and Zeng-Guang Xue analyzed the data; Zhan-Yun Guo planned the experiments and wrote the paper.

## Acknowledgments

We thank some former laboratory members, Ben-Jun Ji, Yu Liu, Ge Song, Lei Zhang, Yu-Qi Guo, Qing-Ping Wu, Li-Li Shou, and Ning Li, for the generation of over 200 GPCR expression constructs. This work was supported by grants from the National Natural Science Foundation of China (grant numbers 31971193, 31470767).

## Appendix A. Supplementary data

Supplementary data to this article can be found online

## Data availability

The data of this study are available in this manuscript, as well as the associated supplementary information.

