## Supplementary Tables and Figures for "CXCL17 activates three MAS-related G protein-coupled receptors independently of its conserved C-terminal fragment"

**Contents**

**Table S1.** Information of the C-terminally LgBiT-fused human MRGPRs generated in this study for the NanoBiT-based β-arrestin recruitment assays.

**Table S2.** Information for generating the expression constructs of human MRGPRs.

**Fig. S1.** Amino acid sequence alignment of human MRGPRs.

**Fig. S2.** Amino acid sequence alignment of some MRGPRX2 orthologs from eutherians.

**Table S1.** Information of the C-terminally LgBiT-fused human MRGPRs generated in this study for the NanoBiT-based β-arrestin recruitment assays. The information of human MRGPRs was retrieved from the gene database of NCBI (https://www.ncbi.nlm.nih.gov/gene). The amino acid sequence of MRGPRs is shown in red, that of LgBiT in blue. The present MRGPRX3 contains an N169D missense SNP (rs4274188) as highlighted in yellow.

| Name | Gene ID | mRNA ID | Protein ID | Amino acid sequence of the C-terminally LgBiT-fused MRGPRs |
|---|---|---|---|---|
| MAS1 | 4142 | NM_002377<br>NM_001366704 | NP_002368<br>NP_001353633 | MDGSNVTSFVVEEPTNISTGRNASVGNAHRQIPIVHWVIMSISPVGFVENGILLWFLCFRMRRNPFTVYITHLSIADISLLFCIFILSIDYALDYELSSGHYYTIVTLSVTFLFGYNTGLYLLTAISVERCLSVLYPIWYRCHRPKYQSALVCALLWALSCLVTTMEYVMCIDREEESHSRNDCRAVIIFIAILSFLVFTPLMLVSSTILVVKIRKNTWASHSSKLYIVIMVTIIIFLIFAMPMRLLYLLYYEYWSTFGNLHHISLLFSTINSSANPFIYFFVGSSKKKRFKESLKVVLTRAFKDEMQPRRQKDNCNTVTVETVVPPVGTGGGSSSGGGMVFTLEDFVGDWEQTAAYNLDQVLEQGGVSSLLQNLAVSVTPIQRIVRSGENALKIDIHVIIPYEGLSADQMAQIEEVFKVVYPVDDHHFKVILPYGTLVIDGVTPNMLNYFGRPYEGIAVFDGKKITVTGTLWNGNKIIDERLITPDGSMLFRVTINS |
| MAS1L | 116511 | NM_052967 | NP_443199 | MVWGKICWFSQRAGWTVFAESQISLSCSLCLHSGDQEAQNPNLVSQLCGVFLQNETNETIHMQMSMAVGQQALPLNIIAPKAVLVSLCGVLLNGTVFWLLCCGATNPYMVYILHLVAADVIYLCCSAVGFLQVTLLTYHGVVFFIPDFLAILSPFSFEVCLCLLVAISTERCVCVLFPIWYRCHRPKYTSNVVCTLIWGLPFCINIVKSLFLTYWKHVKACVIFLKLSGLFHAILSLVMCVSSLTLLIRFLCCSQQQKATRVYAVVQISAPMFLLWALPLSVAPLITDFKMFVTTSYLISLFLIINSSANPIIYFFVGSLRKKRLKESLRVILQRALADKPEVGRNKKAAGIDPMEQPHSTQHVENLLPREHRVDVETPPVGTGGGSSSGGGMVFTLEDFVGDWEQTAAYNLDQVLEQGGVSSLLQNLAVSVTPIQRIVRSGENALKIDIHVIIPYEGLSADQMAQIEEVFKVVYPVDDHHFKVILPYGTLVIDGVTPNMLNYFGRPYEGIAVFDGKKITVTGTLWNGNKIIDERLITPDGSMLFRVTINS |
| MRGPRD | 116512 | NM_198923 | NP_944605 | MNQTLNSSGTVESALNYSRGSTVHTAYLVLSSLAMFTCLCGMAGNSMVIWLLGFRMHRNPFCIYILNLAAADLLFLFSMASTLSLETQPLVNTTDKVHELMKRLMYFAYTVGLSLLTAISTQRCLSVLFPIWFKCHRPRHLSAWVCGLLWTLCLLMNGLTSSFCSKFLKFNEDRCFRVDMVQAALIMGVLTPVMTLSSLTLFVWVRRSSQQWRRQPTRLFVVVLASVLVFLICSLPLSIYWFVLYWLSLPPEMQVLCFSLSRLSSSVSSSANPVIYFLVGSRRSHRLPTRSLGTVLQQALREEPELEGGETPTVGTNEMGAPPVGTGGGSSSGGGMVFTLEDFVGDWEQTAAYNLDQVLEQGGVSSLLQNLAVSVTPIQRIVRSGENALKIDIHVIIPYEGLSADQMAQIEEVFKVVYPVDDHHFKVILPYGTLVIDGVTPNMLNYFGRPYEGIAVFDGKKITVTGTLWNGNKIIDERLITPDGSMLFRVTINS |
| MRGPRE | 116534 | NM_001039165 | NP_001034254 | MMEPREAGQHVGAANGAQEDVAFNLIILSLTEGLGLGGLLGNGAVLWLLSSNVYRNPFAIYLLDVACADLIFLGCHMVAIVPDLLQGRLDFPGFVQTSLATLRFFCYIVGLSLLAAVSVEQCLAALFPAWYSCRRPRHLTTCVCALTWALCLLLHLLLSGACTQFFGEPSRHLCRTLWLVAAVLLALLCCTMCGASLMLLLRVERGPQRPPPRGFPGLILLTVLLFLFCGLPFGIYWLSRNLLWYIPHYFYHFSFLMAAVHCAAKPVVYFCLGSAQGRRLPLRLVLQRALGDEAELGAVRETSRRGLVDIAAPPVGTGGGSSSGGGMVFTLEDFVGDWEQTAAYNLDQVLEQGGVSSLLQNLAVSVTPIQRIVRSGENALKIDIHVIIPYEGLSADQMAQIEEVFKVVYPVDDHHFKVILPYGTLVIDGVTPNMLNYFGRPYEGIAVFDGKKITVTGTLWNGNKIIDERLITPDGSMLFRVTINS |
| MRGPRF | 116535 | NM_145015<br>NM_001098515 | NP_659452<br>NP_001091985 | MAGNCSWEAHPGNRNKMCPGLSEAPELYSRGFLTIEQIAMLPPPAVMNYIFLLLCLCGLVGNGLVLWFFGFSIKRNPFSIYFLHLASADVGYLFSKAVFSILNTGGFLGTFADYIRSVCRVLGLCMFLTGVSLLPAVSAERCASVIFPAWYWRRRPKRLSAVVCALLWVLSLLVTCLHNYFCVFLGRGAPGAACRHMDIFLGILLFLLCCPLMVLPCLALILHVECRARRRQRSAKLNHVILAMVSVFLVSSIYLGIDWFLFWVFQIPAPFPEYVTDLCICINSSAKPIVYFLAGRDKSQRLWEPLRVVFQRALRDGAELGEAGGSTPNTVTMEMQCPPGNASPPVGTGGGSSSGGGMVFTLEDFVGDWEQTAAYNLDQVLEQGGVSSLLQNLAVSVTPIQRIVRSGENALKIDIHVIIPYEGLSADQMAQIEEVFKVVYPVDDHHFKVILPYGTLVIDGVTPNMLNYFGRPYEGIAVFDGKKITVTGTLWNGNKIIDERLITPDGSMLFRVTINS |
| MRGPRG | 386746 | NM_001164377 | NP_001157849 | MFGLFGLWRTFDSVVFYLTLIVGLGGPVGNGLVLWNLGFRIKKGPFSIYLLHLAAADFLFLSCRVGFSVAQAALGAQDTLYFVLTFLWFAVGLWLLAAFSVERCLSDLFPACYQGCRPRHASAVLCALVWTPTLPAVPLPANACGLLRNSACPLVCPRYHVASVTWFLVLARVAWTAGVVLFVWVTCCSTRPRPRLYGIVLGALLLLFFCGLPSVFYWSLQPLLNFLLPVFSPLATLLACVNSSSKPLIYSGLGRQPGKREPLRSVLRRALGEGAELGARGQSLPMGLLPPVGTGGGSSSGGGMVFTLEDFVGDWEQTAAYNLDQVLEQGGVSSLLQNLAVSVTPIQRIVRSGENALKIDIHVIIPYEGLSADQMAQIEEVFKVVYPVDDHHFKVILPYGTLVIDGVTPNMLNYFGRPYEGIAVFDGKKITVTGTLWNGNKIIDERLITPDGSMLFRVTINS |
| MRGPRX1 | 259249 | NM_147199<br>NM_001393578 | NP_671732<br>NP_001380507 | MDPTISTLDTELTPINGTEETLCYKQTLSLTVLTCIVSLVGLTGNAVVLWLLGCRMRRNAFSIYILNLAAADFLFLSGRLIYSLLSFISIPHTISKILYPVMMFSYFAGLSFLSAVSTERCLSVLWPIWYRCHRPTHLSAVVCVLLWALSLLRSILEWMLCGFLFSGADSAWCQTSDFITVAWLIFLCVVLCGSSLVLLIRILCGSRKIPLTRLYVTILLTVLVFLLCGLPFGIQFFLFLWIHVDREVLFCHVHLVSIFLSALNSSANPIIYFFVGSFRQRQNRQNLKLVLQRALQDASEVDEGGGQLPEEILELSGSRLEQPPVGTGGGSSSGGGMVFTLEDFVGDWEQTAAYNLDQVLEQGGVSSLLQNLAVSVTPIQRIVRSGENALKIDIHVIIPYEGLSADQMAQIEEVFKVVYPVDDHHFKVILPYGTLVIDGVTPNMLNYFGRPYEGIAVFDGKKITVTGTLWNGNKIIDERLITPDGSMLFRVTINS |

| | | | | |
|---|---|---|---|---|
| MRGPRX2 | 117194 | NM_054030<br>NM_001303615 | NP_473371<br>NP_001290544 | MDPTTPAWGTESTTVNGNDQALLLLCGKETLIPVFLILFIALVGLVGNGFVLWLLGFRMRRNAFSVYVLSLAGADFLFLCFQIINCLVYLSNFFCSISINFPSFFTTVMTCAYLAGLSMLSTVSTERCLSVLWPIWYRCRRPRHLSAVVCVLLWALSLLLSILEGKFCGFLFSDGDSGWCQTFDFITAAWLIFLFMVLCGSSLALLVRILCGSRGLPLTRLYLTILLTVLVFLLCGLPFGIQWFLILWIWKDSDVLFCHIHPVSVVLSSLNSSANPIIYFFVGSFRKQWRLQQPILKLALQRALQDIAEVDHSEGCFRQGTPEMSRSSLVPPVGTGGGSSSGGGMVFTLEDFVGDWEQTAAYNLDQVLEQGGVSSLLQNLAVSVTPIQRIVRSGENALKIDIHVIIPYEGLSADQMAQIEEVFKVVYPVDDHHFKVILPYGTLVIDGVTPNMLNYFGRPYEGIAVFDGKKITVTGTLWNGNKIIDERLITPDGSMLFRVTINS |
| MRGPRX3 | 117195 | NM_054031<br>NM_001370464 | NP_473372<br>NP_001357393 | MDSTIPVLGTELTPINGREETPCYKQTLSFTGLTCIVSLVALTGNAVVLWLLGCRMRRNAVSIYILNLVAADFLFLSGHIICSPLRLINIRHPISKILSPVMTFPYFIGLSMLSAISTERCLSILWPIWYHCRRPRYLSSVMCVLLWALSLLRSILEWMFCDFLFSGADSVWCETSDFITIAWLVFLCVVLCGSSLVLLVRILCGSRKMPLTRLYVTILLTVLVFLLCGLPFGIQWALFSRIHLDWKVLFCHVHLVSIFLSALNSSANPIIYFFVGSFRQRQNRQNLKLVLQRALQDTPEVDEGGGWLPQETLELSGSRLEQPPVGTGGGSSSGGGMVFTLEDFVGDWEQTAAYNLDQVLEQGGVSSLLQNLAVSVTPIQRIVRSGENALKIDIHVIIPYEGLSADQMAQIEEVFKVVYPVDDHHFKVILPYGTLVIDGVTPNMLNYFGRPYEGIAVFDGKKITVTGTLWNGNKIIDERLITPDGSMLFRVTINS |
| MRGPRX4 | 117196 | NM_054032 | NP_473373 | MDPTVPVFGTKLTPINGREETPCYNQTLSFTVLTCIISLVGLTGNAVVLWLLGYRMRRNAVSIYILNLAAADFLFLSFQIIRLPLRLINISHLIRKILVSVMTFPYFTGLSMLSAISTERCLSVLWPIWYRCRRPTHLSAVVCVLLWGLSLLFSMLEWRFCDFLFSGADSSWCETSDFIPVAWLIFLCVVLCVSSLVLLVRILCGSRKMPLTRLYVTILLTVLVFLLCGLPFGILGALIYRMHLNLEVLYCHVYLVCMSLSSLNSSANPIIYFFVGSFRQRQNRQNLKLVLQRALQDKPEVDKGEGQLPEESLELSGSRLGPPPVGTGGGSSSGGGMVFTLEDFVGDWEQTAAYNLDQVLEQGGVSSLLQNLAVSVTPIQRIVRSGENALKIDIHVIIPYEGLSADQMAQIEEVFKVVYPVDDHHFKVILPYGTLVIDGVTPNMLNYFGRPYEGIAVFDGKKITVTGTLWNGNKIIDERLITPDGSMLFRVTINS |

**Table S2.** Information for generating the expression constructs of human MRGPRs.

| Expression constructs | Vectors for cloning | Restriction enzymes for cleavage of the vector | Oligo primers for PCR amplification of MRGPRs (5' to 3') | Template for PCR amplification | Restriction enzymes for cleavage of the PCR product | Approach for construct generation |
|---|---|---|---|---|---|---|
| pcDNA6/MAS1 | pcDNA6 | NheI; AgeI | Forward: AAG GCT AGC ATG GAT GGG TCA AAC GTG ACA<br>Reverse: AA ACC GGT GG GAC GAC AGT CTC AAC TGT GAC | Human genomic DNA | NheI; AgeI | Ligation by T4 DNA ligase |
| pTRE-BI/MAS1-LgBiT: SmBiT-ARRB2 | pTRE-BI/GPR182-LgBiT: SmBiT-ARRB2 | NheI; AgeI (removing GPR182) | Forward: CCG TCA GAT CGC CTG GAG AAT TCG GGG AGA CCC AAG CTG GCT AGC<br>Reverse: GCT AGA CCC TCC GCC GGT ACC GAC CGG TGG GAC GAC AGT CTC | pcDNA6/MAS1 | No cleavage | Gibson assembly |
| PB-TRE/MAS1 | PB-TRE/ dCas9-VPR | NheI; PmeI (removing dCase9-VPR) | Forward: TTC CTA CCC TCG TAA AGG TCT AGA G CTC ACT ATA GGG AGA CCC AAG CT<br>Reverse: GTT TCA GTT AGC CTC CCC CGT TT C ACA GTC GAG GCT GAT CAG CGG | pcDNA6/MAS1 | No cleavage | Gibson assembly |
| pcDNA6/MAS1L | pcDNA6 | NheI; AgeI | Forward: AGG GCT AGC ATG GTC TGG GGG AAA ATT TGC T<br>Reverse: AA ACC GGT GG TGT TTC CAC ATC GAC CCT GTG C | Human genomic DNA | NheI; AgeI | Ligation by T4 DNA ligase |
| pTRE-BI/MAS1L-LgBiT: SmBiT-ARRB2 | pTRE-BI/GPR182-LgBiT: SmBiT-ARRB2 | NheI; AgeI (removing GPR182) | Forward: CCG TCA GAT CGC CTG GAG AAT TCG GGG AGA CCC AAG CTG GCT AGC<br>Reverse: GCT AGA CCC TCC GCC GGT ACC GAC CGG TGG TGT TTC CAC ATC | pcDNA6/MAS1L | No cleavage | Gibson assembly |
| pcDNA6/MRGPRD | pcDNA6 | NheI; AgeI | Forward: AAG GCT AGC ATG AAC CAG ACT TTG AAT AGC A<br>Reverse: AA ACC GGT GG AGC CCC CAT CTC ATT GGT GCC | Human genomic DNA | NheI; AgeI | Ligation by T4 DNA ligase |
| pTRE-BI/MRGPRD-LgBiT: SmBiT-ARRB2 | pTRE-BI/GPR182-LgBiT: SmBiT-ARRB2 | NheI; AgeI (removing GPR182) | Forward: CCG TCA GAT CGC CTG GAG AAT TCG GGG AGA CCC AAG CTG GCT AGC<br>Reverse: GCT AGA CCC TCC GCC GGT ACC GAC CGG TGG AGC CCC CAT CTC | pcDNA6/MRGPRD | No cleavage | Gibson assembly |
| pcDNA6/MRGPRE | pcDNA6 | NheI; AgeI | Forward: AAG GCT AGC ATG ATG GAG CCC AGA GAA GCT<br>Reverse: AA ACC GGT GG GGC TGC TAT GTC CAC CAG GCC | Human genomic DNA | NheI; AgeI | Ligation by T4 DNA ligase |
| pTRE-BI/MRGPRE-LgBiT: SmBiT-ARRB2 | pTRE-BI/GPR182-LgBiT: SmBiT-ARRB2 | NheI; AgeI (removing GPR182) | Forward: CCG TCA GAT CGC CTG GAG AAT TCG GGG AGA CCC AAG CTG GCT AGC<br>Reverse: GCT AGA CCC TCC GCC GGT ACC GAC CGG TGG GGC TGC TAT GTC | pcDNA6/MRGPRE | No cleavage | Gibson assembly |
| pcDNA6/MRGPRF | pcDNA6 | NheI; AgeI | Forward: AAG GCT AGC ATG GCT GGA AAC TGC TCC TGG GA<br>Reverse: AA ACC GGT GG GGA GGC GTT CCC CGG GGG ACA | Human genomic DNA | NheI; AgeI | Ligation by T4 DNA ligase |
| pTRE-BI/MRGPRF-LgBiT: SmBiT-ARRB2 | pTRE-BI/GPR182-LgBiT: SmBiT-ARRB2 | NheI; AgeI (removing GPR182) | Forward: CCG TCA GAT CGC CTG GAG AAT TCG GGG AGA CCC AAG CTG GCT AGC<br>Reverse: GCT AGA CCC TCC GCC GGT ACC GAC CGG TGG GGA GGC GTT CCC | pcDNA6/MRGPRF | No cleavage | Gibson assembly |
| pcDNA6/MRGPRG | pcDNA6 | NheI; BamHI | Forward: AAG GCT AGC ATG TTT GGG CTG TTC GGC CTC T<br>Reverse: AAG GGA TCC CG TAG GAG ACC CAT GGG CAG GGA CT | Human genomic DNA | NheI; BamHI | Ligation by T4 DNA ligase |
| pTRE-BI/MRGPRG-LgBiT: SmBiT-ARRB2 | pTRE-BI/GPR182-LgBiT: SmBiT-ARRB2 | NheI; AgeI (removing GPR182) | Forward: CCG TCA GAT CGC CTG GAG AAT TCG GGG AGA CCC AAG CTG GCT AGC<br>Reverse: GCT AGA CCC TCC GCC GGT ACC GAC CGG TGG TAG GAG ACC CAT GGG CAG GGA | pcDNA6/MRGPRG | No cleavage | Gibson assembly |
| pcDNA6/MRGPRX1 | pcDNA6 | NheI; AgeI | Forward: AAG GCT AGC ATG GAT CCA ACC ATC TCA ACC TT<br>Reverse: AA ACC GGT GG CTG CTC CAA TCT GCT TCC CGA CA | Human genomic DNA | NheI; AgeI | Ligation by T4 DNA ligase |
| pTRE-BI/MRGPRX1-LgBiT: SmBiT-ARRB2 | pTRE-BI/GPR182-LgBiT: SmBiT-ARRB2 | NheI; AgeI (removing GPR182) | Forward: CCG TCA GAT CGC CTG GAG AAT TCG GGG AGA CCC AAG CTG GCT AGC<br>Reverse: GCT AGA CCC TCC GCC GGT ACC GAC CGG TGG CTG CTC CAA TCT | pcDNA6/MRGPRX1 | No cleavage | Gibson assembly |
| PB-TRE/MRGPRX1 | PB-TRE/ dCas9-VPR | NheI; PmeI (removing dCase9-VPR) | Forward: TTC CTA CCC TCG TAA AGG TCT AGA G CTC ACT ATA GGG AGA CCC AAG CT<br>Reverse: GTT TCA GTT AGC CTC CCC CGT TT C ACA GTC GAG GCT GAT CAG CGG | pcDNA6/MRGPRX1 | No cleavage | Gibson assembly |
| pcDNA6/MRGPRX2 | pcDNA6 | NheI; AgeI | Forward: AAG GCT AGC ATG GAT CCA ACC ACC CCG GCC T<br>Reverse: AA ACC GGT GG CAC CAG ACT GCT TCT CGA CAT | Human genomic DNA | NheI; AgeI | Ligation by T4 DNA ligase |
| pTRE-BI/MRGPRX2-LgBiT: SmBiT-ARRB2 | pTRE-BI/GPR182-LgBiT: SmBiT-ARRB2 | NheI; AgeI (removing GPR182) | Forward: CCG TCA GAT CGC CTG GAG AAT TCG GGG AGA CCC AAG CTG GCT AGC<br>Reverse: GCT AGA CCC TCC GCC GGT ACC GAC CGG TGG CAC CAG ACT GCT | pcDNA6/MRGPRX2 | No cleavage | Gibson assembly |
| PB-TRE/MRGPRX2 | PB-TRE/ dCas9-VPR | NheI; PmeI (removing dCase9-VPR) | Forward: TTC CTA CCC TCG TAA AGG TCT AGA G CTC ACT ATA GGG AGA CCC AAG CT<br>Reverse: GTT TCA GTT AGC CTC CCC CGT TT C ACA GTC GAG GCT GAT CAG CGG | pcDNA6/MRGPRX2 | No cleavage | Gibson assembly |

| | | | | | | |
|---|---|---|---|---|---|---|
| pcDNA6/MRGPRX3 | pcDNA6 | NheI; AgeI | Forward: AAG GCT AGC ATG GAT TCA ACC ATC CCA GTC TT<br>Reverse: AA ACC GGT GG CTG CTC CAA TCT GCT TCC CGA CA | Human genomic DNA | NheI; AgeI | Ligation by T4 DNA ligase |
| pTRE-BI/MRGPRX3-LgBiT:SmBiT-ARRB2 | pTRE-BI/GPR182-LgBiT:SmBiT-ARRB2 | NheI; AgeI (removing GPR182) | Forward: CCG TCA GAT CGC CTG GAG AAT TCG GGG AGA CCC AAG CTG GCT AGC<br>Reverse: GCT AGA CCC TCC GCC GGT ACC GAC CGG TGG CTG CTC CAA TCT | pcDNA6/MRGPRX3 | No cleavage | Gibson assembly |
| pcDNA6/MRGPRX4 | pcDNA6 | NheI; AgeI | Forward: AAG GCT AGC ATG GAT CCA ACC GTC CCA GTC T<br>Reverse: AA ACC GGT GG TGG CCC CAA TCT GCT TCC CGA C | Human genomic DNA | NheI; AgeI | Ligation by T4 DNA ligase |
| pTRE-BI/MRGPRX4-LgBiT:SmBiT-ARRB2 | pTRE-BI/GPR182-LgBiT:SmBiT-ARRB2 | NheI; AgeI (removing GPR182) | Forward: CCG TCA GAT CGC CTG GAG AAT TCG GGG AGA CCC AAG CTG GCT AGC<br>Reverse: GCT AGA CCC TCC GCC GGT ACC GAC CGG TGG TGG CCC CAA TCT | pcDNA6/MRGPRX4 | No cleavage | Gibson assembly |

pTRE3G-BI/GPR182-LgBiT:SmBiT-ARRB2 was generated in our laboratory (unpublished data).

PB-TRE/dCas9-VPR was from Addgene (cat#: 63800).

For oligo primers, the restriction enzyme cleavage site or the fragment pairing with vector is highlighted in yellow, the fragment pairing with PCR template is underlined.

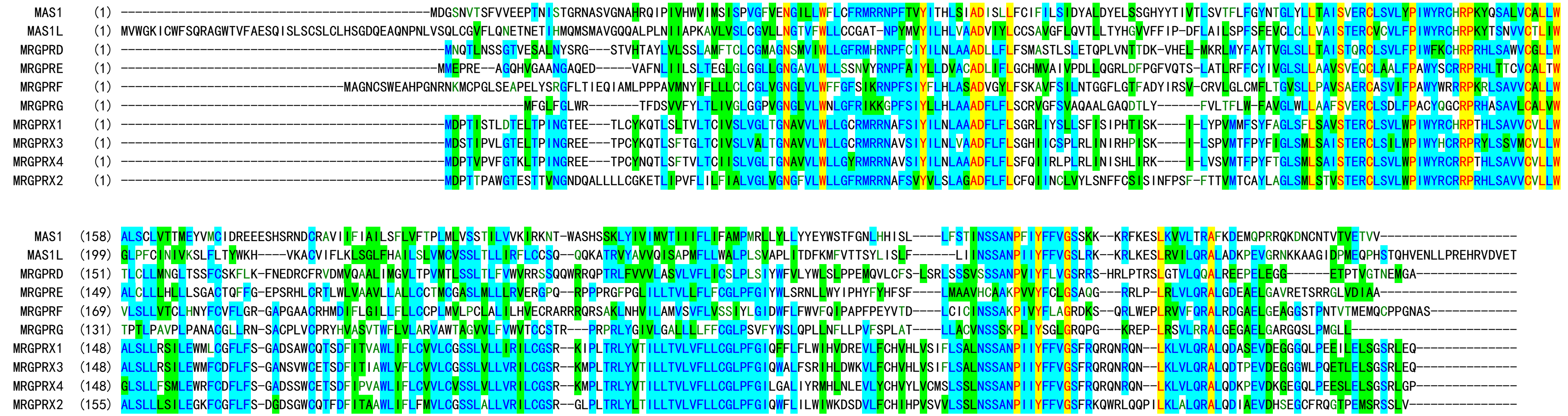


**Fig. S1.** Amino acid sequence alignment of human MRGPRs.

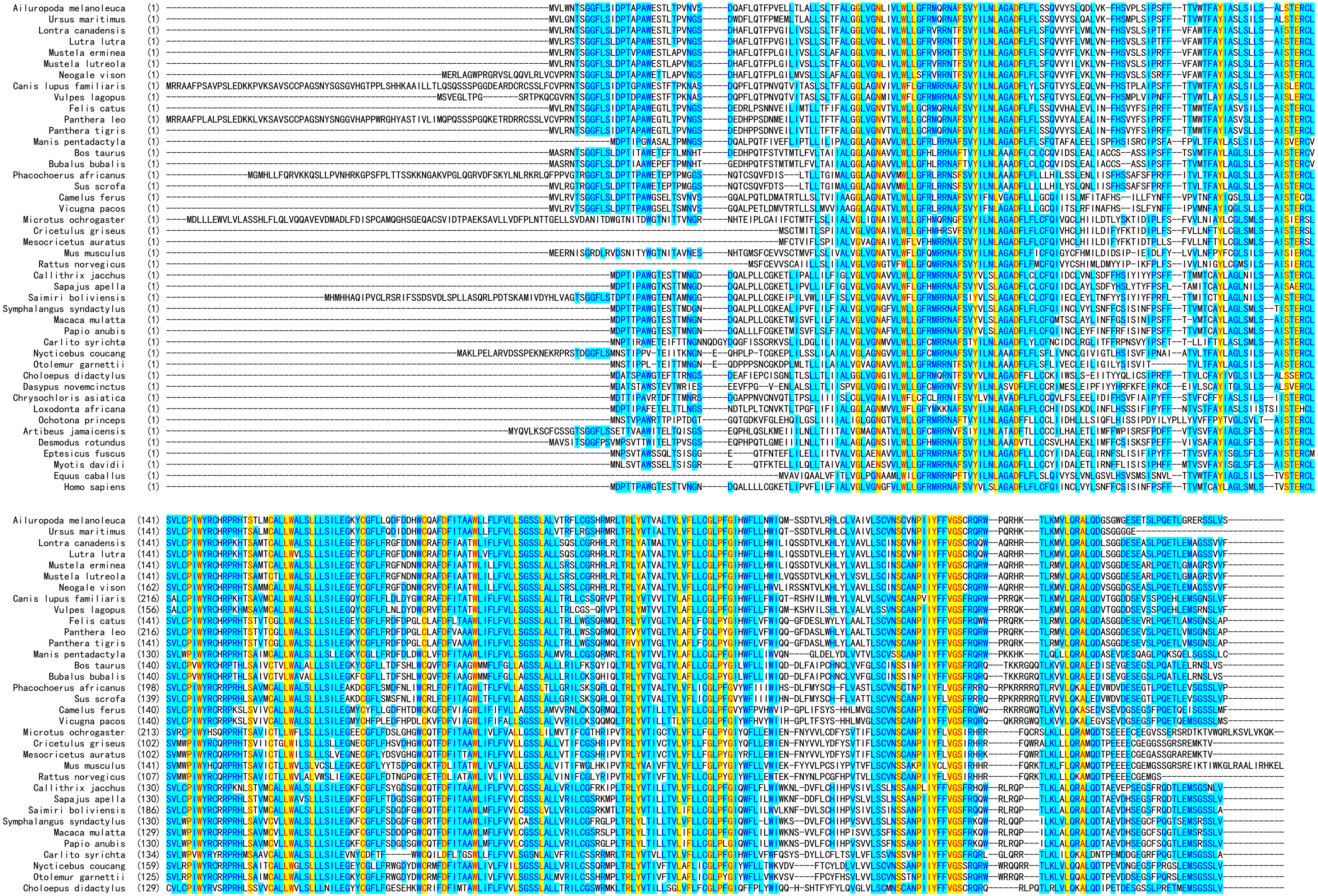
Ailuropoda melanoleuca
Ursus maritimus
Lontra canadensis
Lutra lutra
Mustela erminea
Mustela lutreola
Neogale vison
Canis lupus familiaris
Vulpes lagopus
Felis catus
Panthera leo
Panthera tigris
Manis pentadactyla
Bos taurus
Bubalus bubalis
Phacochoerus africanus
Sus scrofa
Camelus ferus
Vicugna pacos
Microtus ochrogaster
Cricetulus griseus
Mesocricetus auratus
Mus musculus
Rattus norvegicus
Callithrix jacchus
Sapajus apella
Saimiri boliviensis
Symphalangus syndactylus
Macaca mulatta
Papio anubis
Carlito syrichta
Nycticebus coucang
Otolemur garnettii
Choloepus didactylus
Dasypus novemcinctus
Chrysochloris asiatica
Loxodonta africana
Ochotona princeps
Artibeus jamaicensis
Desmodus rotundus
Eptesicus fuscus
Myotis davidii
Equus caballus
Homo sapiens

```
Dasypus novemcinctus    (127) SVLCPVWYRCSRPKHTSSVVCALLWALSLLLSILDGNYCGFLFSDFEDTWCRIFDFITAAWLIFLFAVLSGSSLALLTRMLCGSWRMQLTRPYVAILLSVLAFLLCGLPYGISWFLLFWTLK-HPTLSCSVHCPVGLVLACVNSCANPIIYFFVGSFRHRQ------RLRQTLRLVLQRALQDVPETDDSGASLPPETMKMA-----------------
Chrysochloris asiatica  (130) SILCPVWFRCHRPRHTSATMCALLWVLSLLLSTLEGNYCGFLFEDFEHNWCQTFDFITTAWLIFLFLVLAGCSLVLLNRILYSFQRLQLTRLYVTIVLTVLVFLVCGLPFGIHWFLLIWIH---KNVFTCYLDWVGLILCSVNSSVNPIIYFFVGSFRQRGQLRQRRRLPLTLRIVLQRALQDTPELDESAGSFTQEMSRNSVM---------------
Loxodonta africana      (132) SVLCPIWYHCCHPSHMSVVMCALLWIVSLLLSILEGIYCGFLFRDFKDGWCQTFDFITTAWLMFLFVILSGSSLALMSRILDSSQKMQLTRLYVTIALTVLVFFLYGLPYGISWLLLIWIHS-YLNMFFCSLYIVGVVPSCVNSSANPIIYFFVGSFSQWG--QQRR-FPRTIKTVLQEALQDTPEVDEPGGSLTQEIMKLSGTSVV------------
Ochotona princeps       (132) SVLWPIWYHCHRSKNLSAVVCALLWVLALILNLLECIYCIRLSKHHDDYQCKKIDYTIASWLIFLFVVLLGSSLALLVRVLRGSHRVRLSRLYVTVLLTVLVFLLCGLPFGLFWFLITKNIP-LNFKHLSIVHLIVAVLSGVNSCANPIIYFFVGSFRQQ--------KRQPLKLVLERALQGTPEDDE------------------------------
Artibeus jamaicensis    (150) SITWPIWYRCRRPRHMSAVMCALLWTMSLLLSILQGNYCGFLNRNVHPFWCPALDFITVAWLMLLFVLLSGSSLVLLTRLLCGSQRVQPTRLYVTVGLTVLVFLTCGLPFGIHWFLIFWFQK--DTRAFFRLFLIAVFLSCVNSCANPIIYFFVGSFRQRW--WKRQQ---TLRLVLQRALQDTPEVDKHEGSLPQEILKMSGNSLLS-----------
Desmodus rotundus       (142) SITWPIWYRCRRPRHMSAVTCVLLWTMSLLLSIVEGNYCGFLNRNVHHFWCPALDFITVAWLILLFVLLSGSSLVLLTRLLCGSQRVQPTRLYVTVGLTVLVFLFCGMPFGIHWFLIVWFQE--DIDDFFRLYLIAILLSCLNSCANPVIYFFVGSFRQRW--WKRRQ---TLRLVLQRALQDTPEVDEHGESLPQEPLEMSGSSLVS-----------
Eptesicus fuscus        (126) SVLWPIWYRCRRPRHGSAGMCALLWALSLLLSILEGNYCGFLVRNIHHVWCPVLDFITVAWLILLFVLLSGSSLALMIRMLCGSNRVPPTRLYATIVLTVLVFLLCGLPFGIHWFLLIWLQK--DFDAFVPHFLAAVLLSCVNSCANPIIYFFVGSFRQRW--WKWRH---ALRLILQRALQDTPEVDEPGESLPGETLFMSGS-LVS-----------
Myotis davidii          (126) SVLWPIWYRCRRPRNRSAGMCALLWALSLLLSILEGNYCGFLVRSMHHVWCPVLDFITVAWLTLLFMLLTGSSLALMTRVLCGSNRVPLTRLYATVVLTVLVFFLCGLPYGIIWFLVFWFQK--DLGAFIPHFLAAVLLSCVNSCANPIIYFFVGSFRQRW--WKRRH---TLRLVLQRALQDTPEVDEPGESLSGETLAMSGN-LVS-----------
Equus caballus          (102) SILCPIWYRCRRPRYSSAVICALLWALSLVLSVLEGKYCGFLSRDPHYHWCRVLDLITATWLIFLCVLLSGSSLTLLTRLLCGSQRVPLTRLYVTIVLTVLVFLLCGLPFGMYWFLLIWVYHG--VDVFPFLHLFTTVLSWVNSSANPIIYFFVGSFR-----RRQRGR--TLKLVLQRALQDSPEAEGSEGSLPPETLEMSRSSLVSR----------
Homo sapiens            (130) SVLWPIWYRCRRPRHLSAVVCVLLWALSLLLSILEGKFCGFLFSDGDSGWCQTFDFITAAWLIFLFMVLCGSSLALLVRILCGSRGLPLTRLYLTILLTVLVFLLCGLPFGIQWFLILWIWKDS-DVLFCHIHPVSVVLSSLNSSANPIIYFFVGSFRKQW--RLQQP---ILKLALQRALQDIAEVDHSEGCFRQGTPEMSRSSLV------------
```

**Fig. S2.** Amino acid sequence alignment of some MRGPRX2 orthologs from eutherians.
